# A transcriptomics-guided drug target discovery strategy identifies novel receptor ligands for lung regeneration

**DOI:** 10.1101/2021.05.18.444655

**Authors:** Xinhui Wu, I. Sophie T. Bos, Thomas M. Conlon, Meshal Ansari, Vicky Verschut, Lars A. Verkleij, Angela D’Ambrosi, Aleksey Matveyenko, Herbert B. Schiller, Melanie Königshoff, Martina Schmidt, Loes E. M. Kistemaker, Ali Önder Yildirim, Reinoud Gosens

## Abstract

Currently, there is no pharmacological treatment targeting defective tissue repair in chronic disease. Here we utilized a transcriptomics-guided drug target discovery strategy using gene signatures of smoking-associated chronic obstructive pulmonary disease (COPD) and from mice chronically exposed to cigarette smoke, identifying druggable targets expressed in alveolar epithelial progenitors of which we screened the function in lung organoids. We found several drug targets with regenerative potential of which EP and IP prostanoid receptor ligands had the most significant therapeutic potential in restoring cigarette smoke-induced defects in alveolar epithelial progenitors *in vitro* and *in vivo*. Mechanistically, we discovered by using scRNA-sequencing analysis that circadian clock and cell cycle/apoptosis signaling pathways were enriched in alveolar epithelial progenitor cells in COPD patients and in a relevant model of COPD, which was prevented by PGE2 or PGI2 mimetics. Conclusively, specific targeting of EP and IP receptors offers therapeutic potential for injury to repair in COPD.

## Introduction

One of the main challenges in pharmacology today is the generation of drugs with regenerative potential, with the ability to restore tissue repair in chronic disease. Regenerative medicine has thus far mainly focused on transplantation, tissue engineering approaches, stem- or progenitor cell therapy, or a combination of these^1^. A regenerative pharmacological approach will have considerable additional potential because it can be applied on a relatively large scale. Furthermore, it can be used to halt the disease process in an early stage resulting in real disease-modifying treatment. Also, pharmacological targeting may aid or support other regenerative strategies.

There is a need for regenerative pharmacology in respiratory, cardiovascular, and neurological diseases as well as many other disease areas. In respiratory medicine, chronic obstructive pulmonary disease (COPD) is one of the most common lung diseases with a need for regenerative therapies. The disease is characterized by airflow limitation that is not fully reversible, and which deteriorates progressively. The main difficulty underlying COPD pathogenesis is increased tissue destruction in combination with abnormal tissue repair in susceptible individuals. As current therapies do not modify the course of the disease, developing new therapeutic strategies aiming at regeneration of tissue is necessary.

In affected individuals, there is an increase in alveolar air space associated with destruction of alveolar epithelial cells along with reduced capacity of epithelial progenitors to restore this defect. In the distal lung, alveolar type II cells and alveolar epithelial progenitors harbour stem cell capacity and function to maintain alveolar epithelium^2^. These cells reside in a local tissue microenvironment called the niche, which is composed of supporting cells such as fibroblasts and alveolar macrophages. The niche controls adequate activation of the progenitor cell^1–3^ by means of secreted factors such as WNTs, FGFs, retinoic acid, and many other factors that control stemness, proliferation and differentiation^3^.

Like in many chronic diseases associated with ageing, this local lung microenvironment is insufficiently supportive for lung repair in COPD ^1,4,5^. For example, studies have indicated that an imbalance of canonical and noncanonical WNT signaling results in impaired alveolar regeneration in COPD^4,6^. Moreover, lymphotoxin-β (LTβ), released from CD8+ T-cells in COPD, can negatively interfere with repair. LTβ induces noncanonical NFκB signalling, thereby repressing functional Wnt/β-catenin signalling in the lung^5^. Accordingly, the challenge towards successful regenerative pharmacology in COPD needs to take into consideration the specific hostile microenvironment and the abnormal repair process that stands in the way of adequate regeneration in COPD.

In the present study, we hypothesized that a transcriptomics guided drug target discovery strategy based on gene signatures differentially expressed in COPD and in response to cigarette smoke may be used to identify novel druggable gene targets that are specifically involved in defective lung repair in COPD. Our results show that such a strategy coupled to functional studies in organoids yields novel receptor ligands of which EP and IP prostanoid receptor had the most significant potential in counteracting the negative effects of cigarette smoke on alveolar progenitor cell function.

## Results

### Transcriptomics-guided screening to identify novel targets

We set out to identify novel drug targets that may help restore defective lung repair. To achieve this, we utilized a transcriptomics-guided target discovery strategy (described in Fig. 1a) based on gene signatures of COPD lung tissues ^7^ and of a relevant model of cigarette smoke exposure^8^ to identify differentially regulated druggable genes. We found Reactome pathways related to inflammation such as *Neutrophil degranulation* and *Innate immune system* as well as pathways related to the extracellular matrix such as *Extracellular matrix organization* and *Integrin cell surface interactions* to be enriched in both datasets (Fig. 1b-e). We identified 38 individual target genes that were concordantly up- and 30 individual target genes concordantly downregulated. These genes were filtered through the ‘Drug-gene interactions and potential druggability in the drug gene interaction database’ (DGIdb, http://www.dgidb.org/), which rendered 25 druggable upregulated genes and 16 druggable downregulated genes (Fig. 1f). Genes were further filtered for expression in lung epithelial cells or fibroblasts by consulting the human lung cell atlas (https://asthma.cellgeni.sanger.ac.uk/) and lung map (https://lungmap.net/), which yielded 15 druggable target genes.

**Figure 1.**
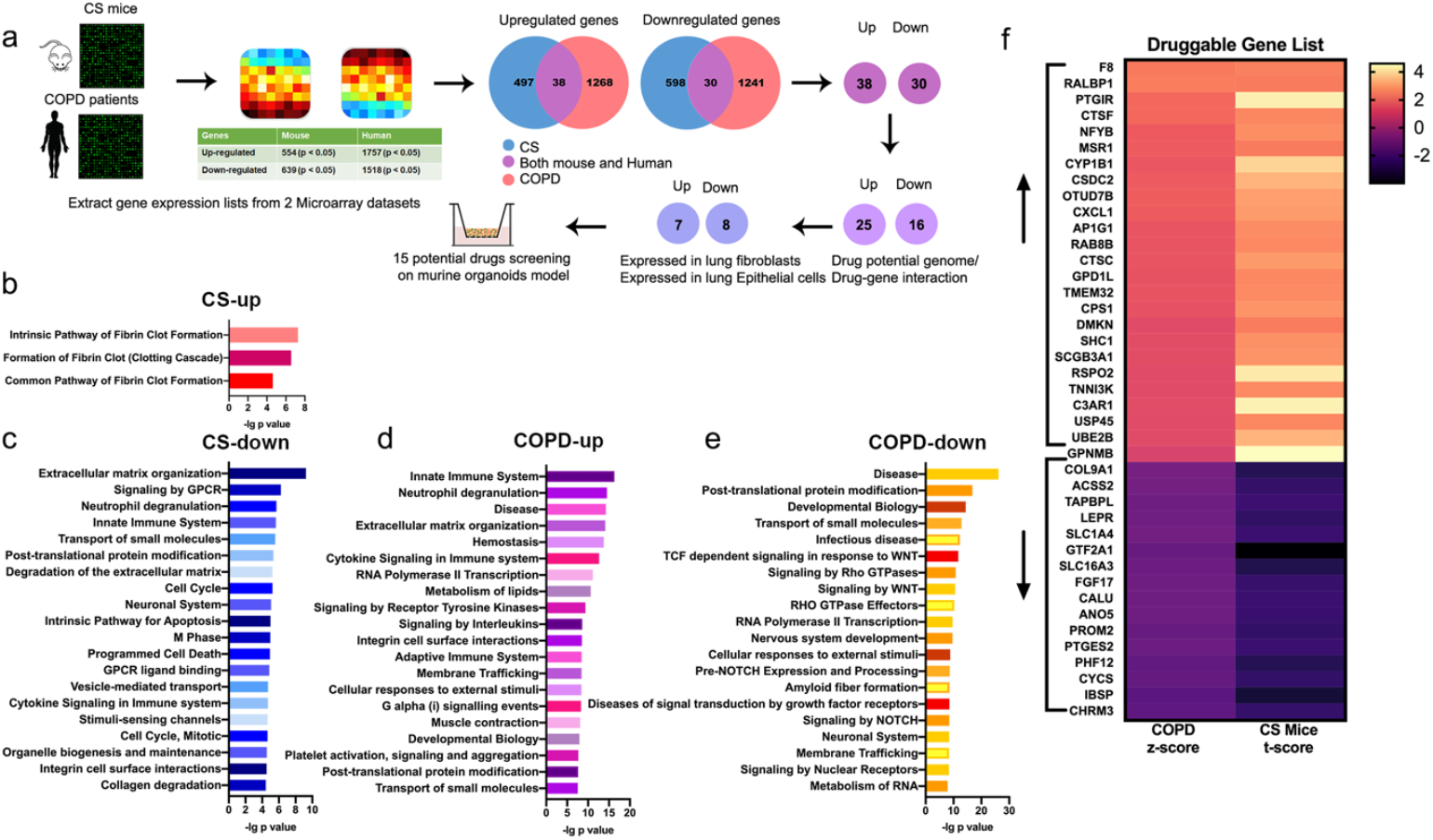
Overview of the transcriptomics-guided drug discovery strategy. **a** Schematic outline of the drug screening strategy. **b-c** Reactome pathway enrichment analysis of differentially up- and downregulated genes from CS-exposed mice^8^ using GSEA (https://www.gsea-msigdb.org/gsea/msigdb/annotate.jsp). **d-e** Reactome pathway enrichment analysis of up- and downregulated genes from COPD patients^7^ using GSEA, the top 20 pathways enriched are presented. **f** Heatmap shows the gene expression pattern of the druggable genes (http://www.dgidb.org/) identified both in CS-exposed mice and COPD patient databases.

To assess the potential relevance of signaling functionally associated with the 15 genes of interest, we set up an *in vitro* organoid model to perform specific drug screening tests. We co-cultured human and mouse CD31-/CD45-/ EpCAM^+^ lung epithelial cells with CCL206 lung fibroblasts in organoids in Matrigel and exposed these *in vitro* to different concentrations (1.25%, 2.5%, 5%) of cigarette smoke extract (CSE) (Fig. 2a, c). The number and size of organoids established by co-culturing human lung tissue derived CD31-/CD45-/EpCAM^+^ cells and MRC5 fibroblasts was significantly decreased by CSE in a concentration dependent manner at day 14 (Fig. 2b). The total number of murine organoids quantified at day 14 of treatment with different concentrations of CSE yielded similar results and was decreased in a CSE dose dependent manner as well (Fig. 2e). To specifically analyze the impact of CSE on organoids derived from alveolar epithelial progenitors, we morphologically subclassified organoids into airway and alveolar type^9^ organoids (Fig. 2d), which revealed that alveolar organoid numbers were more susceptible to cigarette smoke extract exposure than airway organoids (Fig. 2e). Immunofluorescence (IF) studies confirmed that the number of acetylated-α tubulin^+^ (ACT; ciliated cell marker, airway type organoids) and pro-SPC^+^ (type II cell marker, alveolar type organoids) organoids was significantly decreased by 5% CSE (Fig. 2f). The size of both organoid types was decreased at day 14 (Fig. 2g).

**Figure 2.**
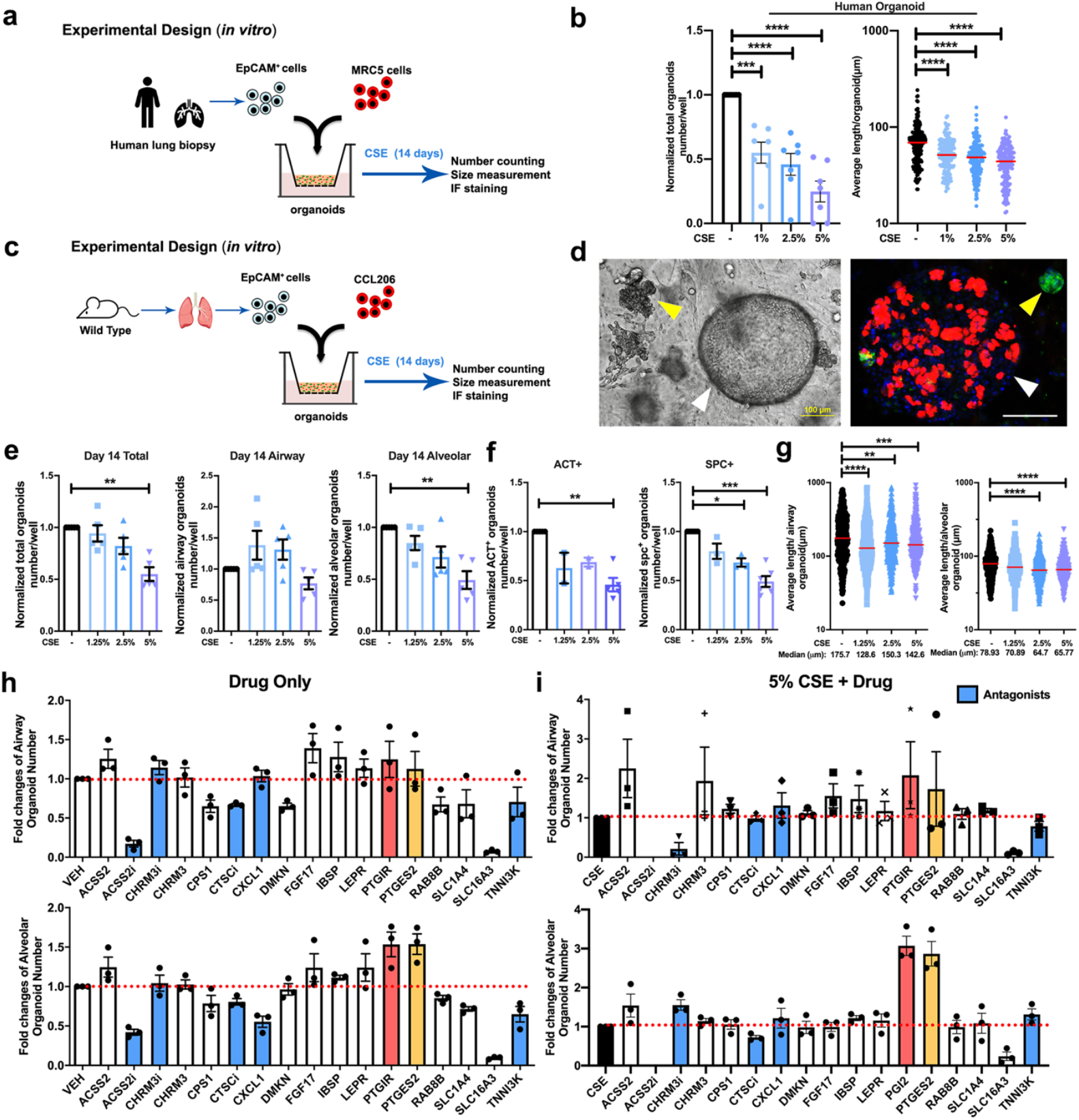
Cigarette smoke exposure represses adult epithelial lung organoid formation. **a** Schematic of *in vitro* human experimental design. **b** Quantification of total amount of human organoids and the quantification of average human organoid diameters after treatment with cigarette smoke extract (CSE) (0, 1, 2.5, and 5%). N = 7 experiments (2 healthy, 5 COPD donors). n > 150 organoids/group. **c** Schematic of *in vitro* murine experimental design. **d** Representative images of murine lung organoids. Left: light microscopy. Right: immunofluorescence of organoids. Green: pro-SPC (SPC), red: acetylated-α tubulin (ACT), blue: DAPI. White arrow: airway type of organoid, yellow arrow: alveolar type of organoid. Scale bar, 100 μm. **e** Quantification of the normalized number of total organoids, airway, and alveolar type organoids on day 14 obtained after treatment with different concentrations of CSE (0, 1.25, 2.5, and 5%). **f** Quantification of normalized ACT^+^ and pro-SPC^+^ organoids obtained after treatment with 0, 1.25, 2.5, and 5% CSE. **g** Quantification of average organoid diameter after treatment with 0, 1.25, 2.5, and 5% CSE measured on day 14. N = 5 experiments, n > 380 organoids/group. **h-i** Overview of drug screening using the *in vitro* murine lung organoid model. Comparison of the normalized number of airway and alveolar type organoids treated with the different drugs of interest in the absence **(h)** or presence **(i)** of 5% CSE. Red bar: PTGIR, yellow bar: PTGES2, blue: antagonists. Data are presented as mean ± SEM in number quantification. Data are presented as scatter plots with medians in size quantification. For all panels: **p < 0.05, **p < 0.01, ***p < 0.001, ****p < 0.0001.

We next aimed to utilize this lung organoid model to evaluate the efficacy of existing COPD therapeutics. Increasing evidence^10,11^ has linked phosphodiesterase (PDE) 4 inhibition to the therapeutic management of respiratory diseases, and roflumilast has been used as an oral medication in COPD patients with a prior history of hospitalization for an acute exacerbation (GOLD 2021). This led us to explore whether the classic PDE4 inhibitor, rolipram, was able to rescue the CS-induced reduction in organoid formation by alveolar progenitors. Thus, organoids were subjected *in vitro* to different concentrations (1 and 10 μM) of rolipram in the presence and absence of 5% CSE for up to 14 days (Fig.S1a). Rolipram (10 μM) alone significantly increased the total number of organoids at day 7 as well as the pro-SPC^+^ organoids at day 14 (Fig.S1b, c), but had no beneficial effects on organoid numbers when combined with CSE exposures. Treatment with rolipram (1 μM) either alone or in combination with CSE induced significantly increase alveolar organoid size (Fig.S1d).

Budesonide is an inhaled corticosteroid used in COPD management^12–15^. Therefore, we examined its effect also in our *in vitro* (Fig. S1e) and *in vivo* (Fig. S1h) CS/organoid models. Intriguingly, budesonide (1-, 10-, 100 nM) in combination with CSE exposure further reduced the number as well as the size of both airway and alveolar type organoids as compared to CSE exposure alone (Fig. S1f, g). *In vivo* exposure to budesonide together with CS increased the number of airway but not alveolar organoids (Fig. S1i), without affecting organoid size (Fig. S1j). Taken together, these data show that *in vitro* exposure to cigarette smoke extract functionally represses human alveolar epithelial progenitor organoid formation, resulting in reduced growth and differentiation, which can be mimicked using murine alveolar epithelial progenitors. Validating the limitations of current pharmacology, PDE4 inhibitors and corticosteroids do not prevent or reduce the detrimental effects of cigarette smoke on organoid formation.

The assay was used subsequently to screen for the functionality of the targets in restoring organoid growth. Genes downregulated in response to CS and COPD were targeted with activating ligands, whereas genes upregulated in response to CS and COPD were targeted using antagonists with exception of *ACSS2* and *CHRM3* for which we included both an agonist and an antagonist. The effects of the drugs targeting the 15 selected genes alone (compared to vehicle, Fig. 2h) and in the presence of 5% CSE exposure (Fig. 2i) on the number of organoids were determined. Specific information of all drug effects on organoid number and size are summarized in Supplementary figure 2 and 3. Interestingly, the compound activating ACSS2 (Acetyl-CoA synthetase short-chain family member 2), increased the number of airway type organoids and the size of alveolar organoids in combination with CSE (Fig. S2-3), whereas the ACSS2 inhibitor had the opposite effects. Atropine (CHRM3 antagonist), IBSP (Integrin Binding Sialoprotien) and LEPR (Leptin receptor) tended to increase the number and size of alveolar organoids in response to CSE as well (Fig. S2-3). However, considering the overall magnitude of alveolar type organoids particularly, PGE2 (target gene *PTGES*) and iloprost, (PGI2 analogue, target gene *PTGIR*) were identified as the by far most promising targets with regards to their capacity in restoring the CSE-induced repression of organoid formation (Fig. 2h-i, S2-3).

### PGE2 and PGI2 significantly prevent alveolar epithelial dysfunction

The *PTGES2* and *PTGIR* genes encode membrane-associated prostaglandin E synthase and the prostacyclin (PGI2) receptor, respectively. PGE2 acts on 4 receptor subtypes, being *PTGER1-4*, whereas PGI2 acts primarily on *PTGIR*. We assessed their expression in human lung tissue of healthy smokers and COPD patients and found maintained expression of all receptors in disease with some small differences in expression, most notably a reduced expression of *PTGER2* and increased expression of *PTGIR* (Fig. 3a). Single cell RNAseq (sc-RNAseq) data from human lung tissue shows similar expression of all 5 receptors was detected in alveolar epithelial cells and in fibroblasts (http://www.copdcellatlas.com/). Single cell RNA sequencing of mouse lung tissue showing that expression of *Ptger2* and *Ptger4* were highest compared to that of *Ptger1* and *Ptger3* in mesenchymal cells (Fig. 3b, c). Interestingly, the expression of *PTGES*, and *PTGES2*, the enzymes responsible for PGE2 synthesis were relatively ubiquitous in human and mouse lung tissue, whereas *PTGIS*, the enzyme responsible for PGI2 synthesis was highest in mesenchymal cell types including fibroblasts. The expression of all these receptors showed lower copy numbers, which is expected for G protein-coupled receptors (GPCRs).

**Figure 3.**
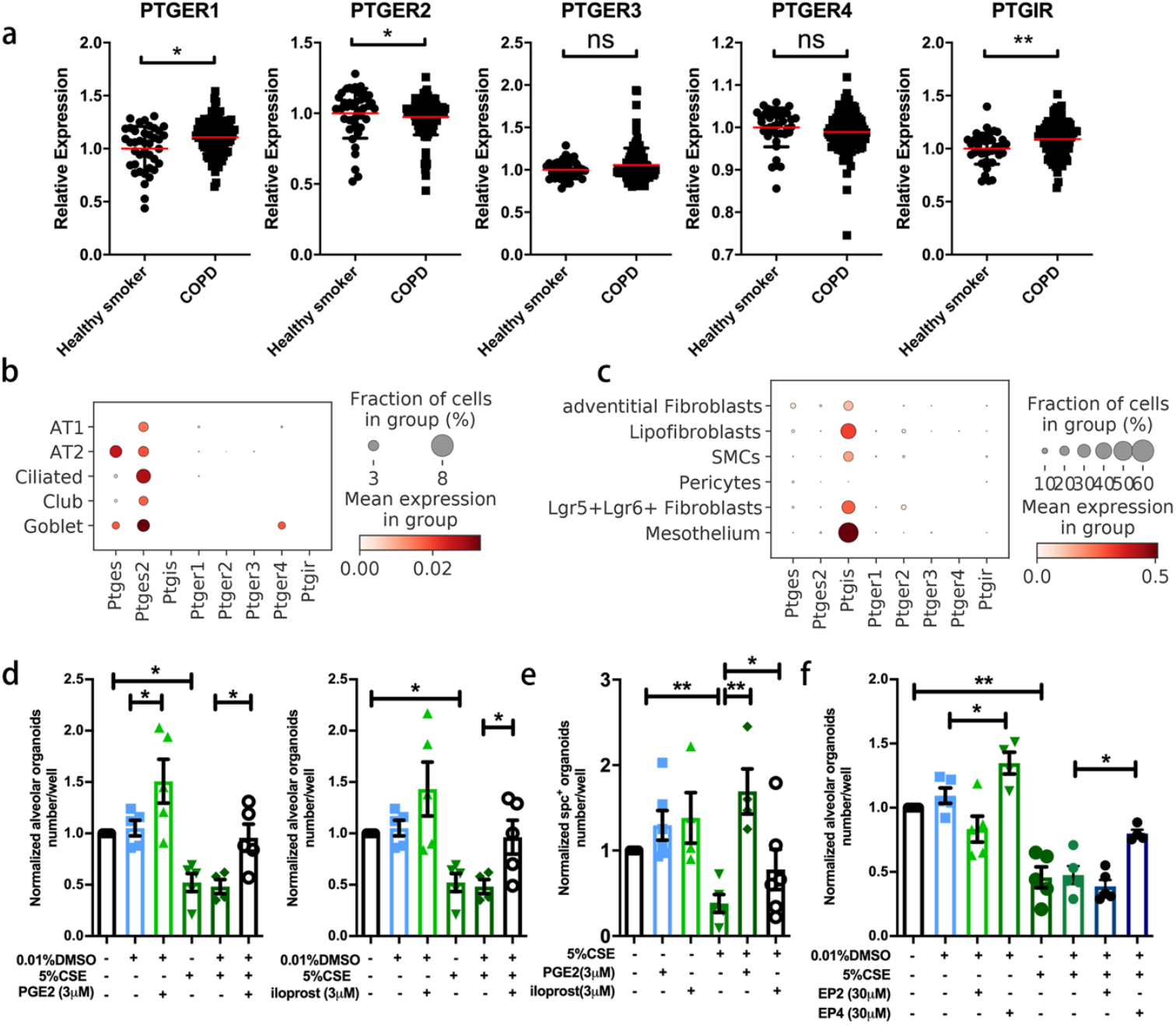
16,16-dimethyl prostaglandin E2, iloprost, and selective EP2 and EP4 analogues restore lung organoid formation in response to cigarette smoke (extract). **a** The relative gene expression of *PTGER1, PTGER2, PTGER3, PTGER4* and *PTGIR* in healthy smokers (N = 40) and COPD patients (N = 111) downloaded from the NCBI GEO database GSE76925. **b-c** Data are extracted from the NCBI GEO database GSE151674 **b** The expression *of Ptges, Ptges2, Ptgis, Ptger1, Ptger2, Ptger3, Ptger4* and *Ptgir* in epithelial cells using scRNA-seq analysis of mouse lung tissue. **c** The expression *of Ptges, Ptges2, Ptgis, Ptger1, Ptger2, Ptger3, Ptger4* and *Ptgir* in mesenchymal cells using scRNA-seq analysis of mouse lung tissue. **d** Quantification of normalized number of alveolar type organoids treated with vehicle control, or 5% CSE ± PGE2 agonist (16,16-dimethyl prostaglandin E2)/iloprost. **e** Quantification of normalized number of SPC^+^ organoids treated with vehicle control, or 5% CSE ± PGE2 agonist (16,16-dimethyl prostaglandin E2) or iloprost. **f** Quantification of normalized number of alveolar type of organoids treated with vehicle control, or 5% CSE ± selective EP2 or EP4 agonist. Data are presented as mean ± SEM. *p < 0.05, **p < 0.01, ***p < 0.001, ****p < 0.0001.

To further characterize the effects of PGE2 and PGI2 on defective alveolar epithelial progenitors, we examined them *in vitro* (Fig. 3d-f) and *in vivo* (Fig. 4) CS(E). The PGE2 analogue 16,16-dimethyl prostaglandin E2 and the prostacyclin analogue iloprost both increased the number of alveolar type organoids even in the presence of 5% CSE (Fig. 3d) and significantly increased the number of SPC^+^ organoids under conditions of CSE exposure (Fig. 3e). To address the relative roles of the two Gs-coupled PGE2 receptors, EP2 and EP4, we evaluated the selective agonists ((R)-Butaprost and 5-[(3S)-3-hydroxy-4-phenyl-1-buten-1-yl]1-[6-(2H-tetrazol-5R-yl)hexyl]-2-pyrrolidinone, CAY10598) of these receptors. Focus was on these Gs coupled receptors, as we found that cholera toxin, a well-known inducer of constitutive adenylyl cyclase activity and cAMP signaling, increased organoid formation both with and without the exposure to CSE (Fig. S4a). The EP2 selective butaprost had no effects on organoid number (Fig. 3f) but increased the alveolar size in the absence and presence of 5% CSE (Fig. S5c). The EP4 selective agonist increased the number of alveolar type organoids significantly and prevented the number reduction resulting from 5% CSE exposure (Fig. 3f). The EP4 agonist also increased the size of both types of organoids in the absence and presence of 5% CSE (Fig. S5c). To explore whether duration of drug exposure affected organoid formation, the organoids were treated *in vitro* with PGE2 analogue or PGI2 analogue for 3 different time windows during organoid development as illustrated in Supplementary figure 5d-g. These time windows were identified previously^9^, and mark the initial division phase (day 0-2), proliferation (day 2-7), and differentiation phase (day 7-14). We observed no effects on the number of organoids for any of the short-term drug treatments, suggesting continuous treatment with or iloprost during all phases of organoid formation is required.

**Figure 4.**
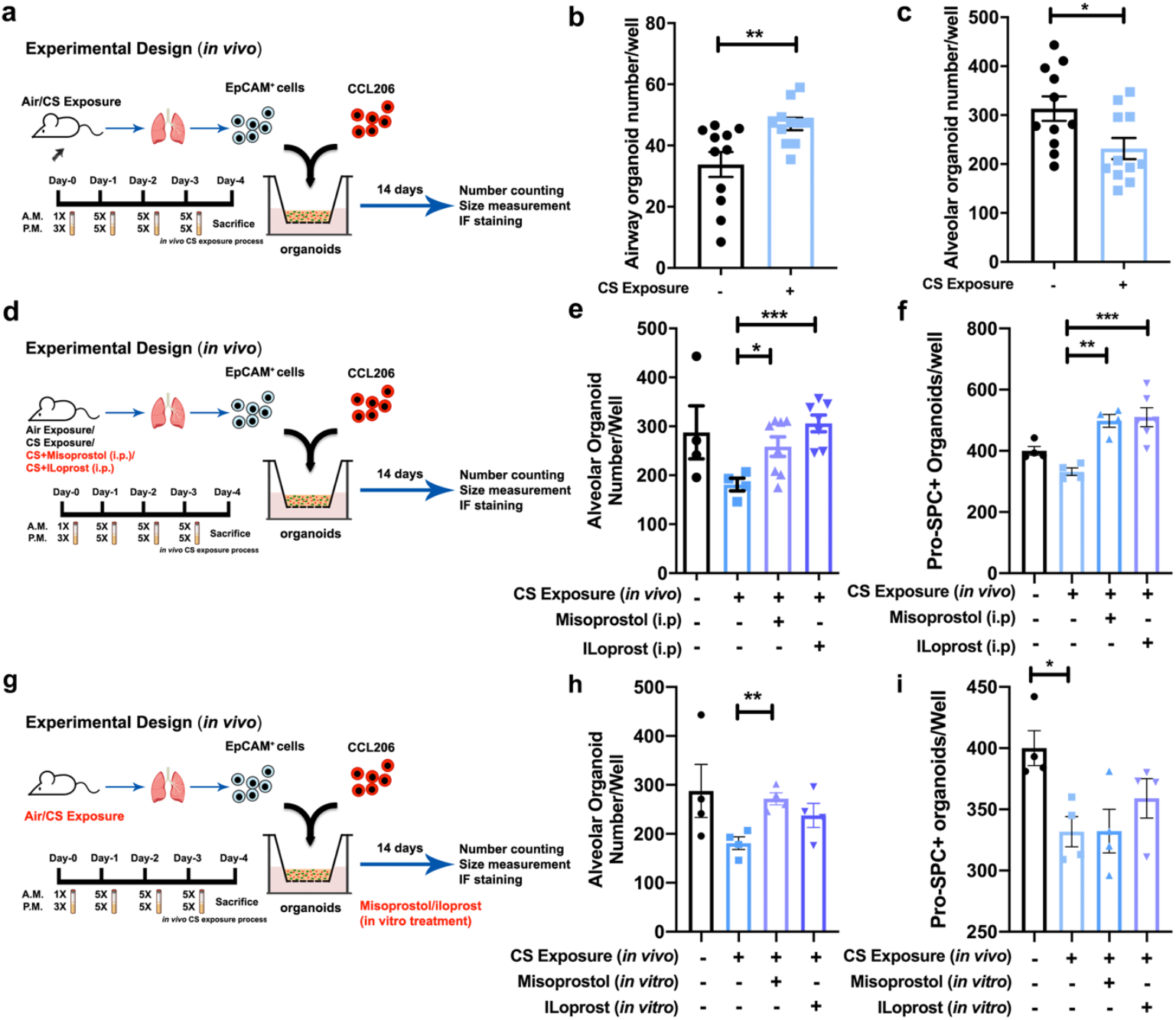
Administration (*in vivo* and *in vitro)* of misoprostol and iloprost to cigarette smoke-exposed mice restored lung organoid formation. **a** Schematic of *in vivo* CS exposure experimental setup. **b-c** Number of airway and alveolar type organoids quantified on day 14 of co-culturing CCL-206 fibroblasts and EpCAM^+^ cells (isolated from air exposed/CS exposed mice). N = 11 experiments. **d** Schematic of experimental design. Organoids were generated from air- or CS-exposed mice treated *in vivo* with misoprostol (i.p.) or iloprost (i.p.); all organoids were treated with normal organoid medium. **e-f** Number of alveolar type and pro-SPC^+^ organoids quantified on day 14 from co-culture of CCL-206 fibroblasts and EpCAM^+^ cells (isolated from air-(control) and CS-exposed mice treated intraperitoneally with misoprostol or iloprost). **g** Schematic of experimental design. Organoids were generated from mice exposed to air or CS. Misoprostol and iloprost were added *in vitro* to the organoid medium for treatment. **h-i** Number of alveolar type and SPC^+^ organoids quantified on day 14 from co-culture of CCL-206 fibroblasts and EpCAM^+^ cells (isolated from air- and CS-exposed mice) treated with misoprostol/iloprost *in vitro*. Data are presented as mean ± SEM. *p < 0.05, **p < 0.01, ***p < 0.001, ****p < 0.0001.

To examine the effects of PGE2 and PGI2 *in vivo*, we exposed mice to air (vehicle control), CS, CS + misoprostol (PGE2 analogue), or CS + iloprost as shown in Figure 4. To assess the impact of *in vivo* cigarette smoke (CS) exposure, we isolated CD31-/CD45-/EpCAM^+^ cells from mice exposed to air or CS and co-cultured these with CCL206 fibroblasts *in vitro* for 14 days (Fig. 4a). Interestingly, the number of alveolar organoids was significantly decreased after *in vivo* CS exposure indicating that a relatively short exposure to cigarette smoke *in vivo* is sufficient to capture early changes in progenitor cell function (Fig. 4c). Cigarette smoke exposure did not change the yield of CD31-/CD45-/EpCAM^+^ cells, whereas exposing mice to misoprostol or iloprost increased the yield of EpCAM+ cells (Fig. S6a). The organoid assay revealed that *in vivo* (i.p.) treatment with either misoprostol or iloprost significantly increased the number of alveolar type and SPC^+^ organoids (Fig. 4e-f) *ex vivo*. Next, to investigate whether *in vitro* drug treatment would have similar effects on damage caused by *in vivo* CS exposure, we isolated EpCAM^+^ cells from either air- or CS-exposed mice and subjected these to *in vitro* misoprostol or iloprost treatment in the organoid assay for 14 days (Fig. 4g). *In vitro* misoprostol increased the number of alveolar type organoids in cultures derived from CS-exposed mice (Fig. 4h). Only *in vitro* misoprostol increased the size of alveolar organoids derived from CS-exposed animals (Fig. S6c). Taken together, our data show that PGE2 and PGI2 analogues protect alveolar epithelial progenitor function from the effects of CS exposure. In addition, EP4 rather than EP2 seems to mediate the protective effects of PGE2.

### Distinct genetic signatures in regulation of defective alveolar epithelial repair

To unravel the transcriptional changes leading to impaired lung organoid formation after exposure to CS as well as the mechanisms underlying the beneficial effects of PGE2 and PGI2 treatment, we performed RNA sequencing (Fig. 5a) on EpCAM^+^ cells isolated from mice exposed to air (control), CS, CS + misoprostol(i.p.), or CS + iloprost (i.p.), directly after the isolation procedure (i.e., prior to inclusion in the organoid assay). Principal-component analysis (PCA) revealed that the CS-exposed group is transcriptionally distinct from the control group (Fig. 5b), and that the CS/misoprostol and CS/iloprost groups are transcriptionally different from the CS exposed group. The top 20 differentially expressed genes from these three comparisons, including both up- and downregulated genes, are shown in the heatmaps (Fig. 5c) and summarized in supplementary materials. Reactome pathway analysis was used to identify molecular pathways overrepresented in CS/misoprostol/iloprost-modulated genes in alveolar epithelial cells. Within the top 20 enriched pathways (Fig. 5d, supplementary materials), genes associated with cell cycle, mitotic prometaphase, DNA replication/synthesis and RHO GTPases activate formins signaling pathways were upregulated by CS exposure compared to air (control) exposure, however, these were downregulated by treatment with misoprostol or iloprost. Notably, genes associated with the circadian clock signaling pathway were downregulated by CS exposure but restored by treatment with misoprostol or iloprost. Moreover, signaling by FGFR1, 3 and 4 were downregulated in response to CS exposure, whereas the same signaling pathways were upregulated by misoprostol but not iloprost treatment. These findings suggest that the repairing mechanisms of misoprostol and iloprost in response to CS were common in cell cycle and circadian clock signaling and the distorted FGFR signaling resulted from CS was corrected by misoprostol uniquely. In addition, the nuclear receptor transcription pathway, growth hormone receptor signaling, MAPK, WNT and cell-cell communication signaling pathways appear to downregulate in response to CS, which were not observed in either misoprostol/iloprost treatment group. Overall, the RNA-seq analysis demonstrates both common and distinct transcriptomic mechanisms of misoprostol and iloprost treatment in response to CS exposure in alveolar epithelial progenitors.

**Figure 5.**
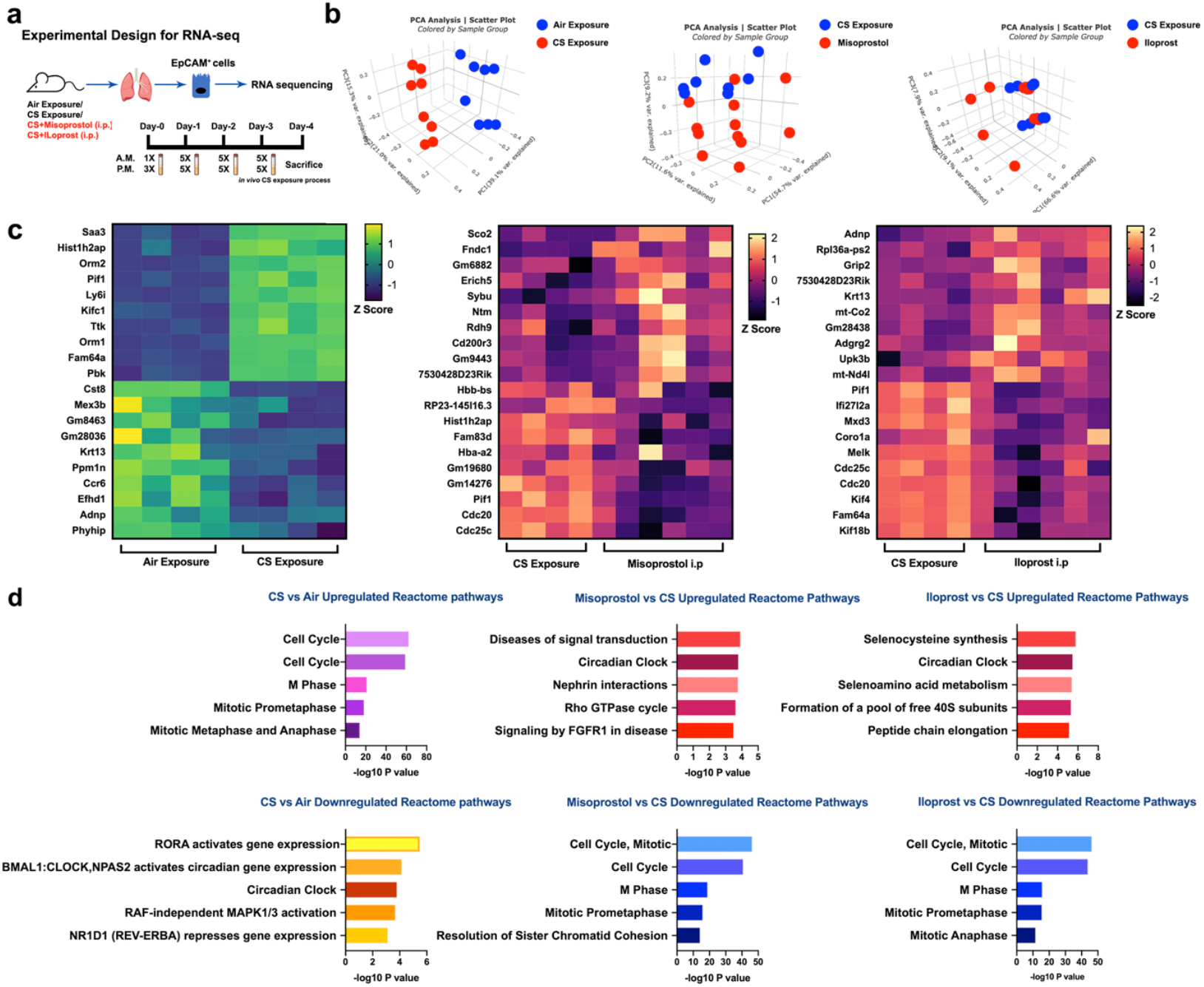
Transcriptomic signatures in response to cigarette smoke with(out) misoprostol and iloprost. **a** Schematic experimental design for RNA sequencing. **b** PCA plots demonstrate the clusters betweern different comparisons: air vs CS, CS vs CS+misoprostol, and CS vs CS+iloprost. **c** Heatmaps displaying the top 20 differentially expressed genes in air vs CS, CS vs CS+misoprostol, and CS+iloprost exposed epithelial cells. **d** The top 5 significantly up- and downregulated reactome pathways enrichment form differentially expressed genes within the comparisons of air vs CS exposure, CS exposure vs CS+misoprostol, and CS exposure vs CS+iloprost.

## Discussion

Chronic obstructive pulmonary disease results from repeated micro-injuries to the epithelium often caused by tobacco smoking. In susceptible individuals, this results in tissue remodeling in the conducting airways and destruction of the respiratory bronchioles and alveoli^19^. No clinically approved pharmacological treatment prevents or reverses the tissue destruction in the distal lung. The results of our study are in line with this contention and demonstrate that the PDE4 inhibitor rolipram and the corticosteroid budesonide had no, or only very limited, beneficial effects on impaired organoid growth and differentiation in response to cigarette smoke extract *in vitro* or in response to cigarette smoke exposure *in vivo*. In fact, if anything, budesonide appeared to restrict progenitor cell growth, which is a concern given the wide use of corticosteroids in the management of COPD. This underscores the need for novel drug targets.

Consequently, we set out to search for new potential drug targets for lung repair in COPD and identify EP and IP receptor agonists as two such potential targets, by utilizing a transcriptomics-guided drug discovery strategy. Both EP and IP receptor agonists were able to promote epithelial repair responses after exposure to cigarette smoke (extract). Whereas PGE2 and PGI2 showed the most profound changes, other methods including ACSS2 agonism, LEPR agonism and IBSP agonism yielded smaller effects. ACSS2 supports acetyl-CoA synthesis from acetate in the cytosol^20,21^, and thereby plays an important role in lipid metabolism and in the regulation of histone acetylation in the nucleus during gene transcription. IBSP is a member of the small integrin-binding ligand N-liked glycoprotein (SIBLING) family^22,23^, which is associated with bone metastases of lung cancer^24^. LEPR (leptin receptor), is an adipocytokine that has a central role in regulating food intake and energy expenditure^25^, but has also been linked to lung function decline in a population in COPD^26^. Nonetheless, among all the candidate targets, PGE2 (*PTGES2*) and PGI2 (*PTGIR*) analogues emerged as the most promising compounds among all drugs in the current study. Prostaglandins (PGs) are lipid mediators synthesized from arachidonic acid (AA) via the cyclooxygenase pathway, and include PGD2, PGI2, PGH2, PGE2 and PGF2α^27^. PGI2 signals via IP receptors to induce cAMP signaling, similar to EP4 receptors. We show EP4 receptors to have similar expression as in non-COPD controls, whereas IP receptors are expressed at higher levels in COPD patients indicating the expression of both receptors is maintained in disease.

We show that PGE2 agonists are beneficial in reducing CS-induced damage in alveolar epithelial progenitors. However, PGE2 has been reported as an unstable molecule with an extremely short half-life; therefore, targeting its receptors with specific more stable analogues may be a better alternative. PGE2 is the most widely produced PG in the human body and it signals via four specific G-protein-coupled receptors (EP1-4^28,29^). The interactions between PGE2 and EP receptors depend on tissue and cell type, specific receptor expression, and differences in binding affinities, leading to unique patterns of EP receptor activation^30^. PGE2 can stimulate cAMP production through EP2 and EP4 receptors, whereas EP3 activation results in decreased cAMP synthesis and EP1 stimulation is coupled to Gq-activation and (enhanced) Ca^2+^ signaling^27,30,31^. EP1 and EP3 receptors can mediate bronchoconstriction indirectly through activation of neural pathways^32^, as a consequence non-selective PGE2 analogues are unsuitable as pulmonary drugs.

Therefore, we selected analogues of EP2 and EP4 to mimic effect of PGE2 in our organoid assay, and demonstrated that EP4 agonism showed beneficial effects against impaired organoid formation in response to (CS)E exposure. Targeting EP4 receptors is worthwhile investigating in more detail in the future, as the effects may surpass epithelial repair only. Additional beneficial effects of EP4 agonism in COPD may include bronchoprotection^33^ and inhibition of inflammation^34^, suggesting that EP4 agonism could unify several functional features that support the treatment of COPD, making this an intriguing pharmacological target. Iloprost, a stable PGI2 analogue ^27,35–39^, has been shown to have anti-inflammatory effects and protects against bleomycin-induced pulmonary fibrosis in mice^36^, and is also clinically used for the treatment of pulmonary hypertension^35^. Although a recent study^38^ showed iloprost improved clinical outcomes in COPD patients with poor lung oxygenation, its impact on alveolar repair is unknown. Here we show that iloprost prevents the repressed organoid formation resulting from CS(E) exposure.

By generating transcriptomic signatures of epithelial progenitors derived from mice exposed *in vivo* to air, CS, CS + misoprostol or CS + iloprost, we uncovered dynamic molecular signaling pathways in response to CS exposure. Intriguingly, we identified circadian clock signaling as being significantly repressed in the alveolar epithelial progenitors derived from mice exposed to CS, which could be improved by either misoprostol or iloprost treatment. Circadian rhythms^40–43^ are normally generated and regulated by clock genes, including *BMAL1* (*ARNTL1*) and *CLOCK* encoding activators, period (*PER1-3*) and cryptochrome genes (*CRY1-2*) that encode repressors, and the nuclear receptors Rev-erb (*NR1D1* and *NR1D2*) and *RORA* which constitute secondary regulatory loops. These core clock genes not only activate or repress a cell-autonomous clock, but also regulate the clock-controlled genes (CCGs)^44^, thus interacting with other molecular signaling pathways. Previously, it has been demonstrated that clock signaling is downregulated in CS exposed mice, linked to an impairment of anti-oxidant defense mechanisms^45^ and Rev-erbα has been shown as an key regulator of inflammatory response in lung injury models^16–18,46^

Furthermore, we found that CS exposure upregulated pathways associated with cell cycle activity in alveolar epithelial progenitors, which could be counteracted by *in vivo* misoprostol or iloprost treatment. The cell cycle^47–51^ is driven by a set of tightly regulated molecular events controlling DNA replication and mitosis with four phases, and each individual cell may require different triggers in order to decide whether to enter proliferation or apoptosis. To further assess alveolar epithelial progenitors under which cell cycle/apoptotic status in response to CS exposure as well as additional PGE2/PGI2 treatments may be the next step to investigate in the future. A link between circadian clock and cell cycle signaling pathways has been proposed^44,50,52^. Importantly, the molecular control of the biological clock is dependent on cAMP signaling and cAMP activators are known to entrain the biological clock^53^, explaining the link between PGE2 and PGI2 activation and restoration of the defective clock signaling in combination with CS exposure. Hence, it is of great interest to determine in more molecular detail how these two oscillatory systems communicate in regulating PGE2/PGI2-mediated lung repair in future studies.

In conclusion, in this study we demonstrate for the first time the protective effects of several drug candidates, and in particular PGE2 and PGI2 analogues, against *in vivo* and *in vitro* CS(E)-induced damage of alveolar epithelial progenitors. Furthermore, using transcriptome analysis, we show that CS induces a wide range of transcriptional changes, including alterations of circadian clock and cell cycle signaling pathways, which can be counteracted by either misoprostol (PGE2) or iloprost (PGI2) treatment. Overall, these data provide promising therapeutic strategies to specifically address defective lung repair in respiratory diseases, in particular targeting EP4 and IP receptors.

## Methods

### Animals

All mouse experiments for organoid study were performed at the Central Animal Facility (CDP) of the University Medical Center Groningen (UMCG) in accordance with the national guidelines and upon approval of the experimental procedures by CDP and the Institutional Animal Care and Use Committee (IACUC) of the University of Groningen (CCD license AVD105002015303). C57BL/6J (555) and BALB/cByJ (Jax-strain) mice (both genders, 8–12 weeks of age) were maintained under 12-h light/ dark cycles and were allowed food and water ad libitum. Animals for circadian clock studies were exposed to CS and/or administrated with compounds at the same time of the day for all mice in all groups. Animals were euthanized at the same time of the day. Adult (female, 8-10 weeks of age) C57BL/6N mice were obtained from Charles River (Sulzfeld, Germany) for the single cell RNAseq analysis of lungs following exposure to chronic CS. These experiments were performed at the Helmholtz Zentrum München and approved by the ethics committee for animal welfare of the local government for the administrative region of Upper Bavaria (Regierungspräsidium Oberbayern) and were conducted under strict governmental and international guidelines in accordance with EU Directive 2010/63/EU.

### Human material

The human lung tissue was obtained from lung transplant donors according to the Eurotransplant guidelines including the absence of primary lung diseases such as asthma and COPD, and no more than 20 pack years of smoking history^54^. Gene expression in human lung published data sets was obtained by down loading series matrix files from the NCBI GEO database for GSE76925^55^ and gene expression normalized to healthy smokers.

### *In vivo* cigarette smoke exposure

Mice (n = 4–11/group, 10-12 weeks old) were exposed (whole body) to 3R4F research cigarettes (Tobacco Research Institute, University of Kentucky, Lexington, KY) for four consecutive days (two sessions/day, 8 hours between each exposure) to establish an acute smoke-induced inflammation model, as described previously^8^. In the cigarette smoke (CS) group, mice were exposed to 1 cigarette in the morning and 3 in the afternoon on day 1. From day 2 to 4, mice were exposed to 5 cigarettes each session. All cigarettes were smoked without a filter in 5 min at a rate of 5L/h in a ratio with 60 L/h air using a peristaltic pump (45 rpm, Watson Marlow 323 E/D, Rotterdam, NL). In the control group, mice were exposed to fresh air using similar exposure chambers as the CS group.

In some studies, budesonide was nebulized (0.1 mM, 15 min/mouse/exposure) to wild-type mice (n = 6) prior to each CS exposure. In separate studies, intraperitoneal (IP) injections of 50 μg misoprostol or 50 μg iloprost were given 30 min to wild-type mice (n = 6-8) prior to each CS exposure. On day 5, mice were sacrificed, and the lungs were immediately used for establishing organoid cultures or stored at - 80°C for further experimental uses.

For the single cell RNAseq analysis of lungs CS was generated from 3R4F Research Cigarettes with the filters removed. Mice were whole body exposed to active 100% mainstream CS of 500 mg/m^3^ total particulate matter (TPM) for 50 min twice per day for 4m in a manner mimicking natural human smoking habits as previously described^55^

### Fibroblast culture

Mouse fibroblasts, CCL206 (Mlg [CCL206], ATCC, Wesel, Germany) were cultured in DMEM/F12 medium supplemented with 10% (v/v) fetal bovine serum, 100 U/mL penicillin/streptomycin, 2 mM L-glutamine, and 1% amphotericin B in a humidified atmosphere under 5% CO2/95% air at 37°C, as previously described^6,9,56^. For organoid experiments, fibroblasts were proliferation-inactivated by incubation in mitomycin C (10 μg/mL, Sigma, M4287) for 2 h, followed by 3 washes with PBS after which the cells were trypsinized before introduction into the organoid co-cultures. Human lung fibroblasts MRC5 (CCL-171; ATCC, Wesel, Germany) were cultured in Ham’s F12 medium supplemented with the same additives as the murine fibroblasts medium.

### Cigarette smoke extract (CSE)

The smoke of two 3R4F research cigarettes was pumped into 25 mL warm fibroblasts culture medium to produce 100% cigarette smoke extract (CSE)^11^. All cigarettes were without a filter and smoke was passed through the medium using a peristaltic pump (45 rpm, Watson Marlow 323 E/D, Rotterdam, NL). CSE was freshly prepared before each set of experiments.

### Organoid culture

The organoid culture system is based on previously published protocols from our group^6,9,56^. In brief, epithelial cells (CD31^-^/CD45^-^/CD326^+^) were freshly isolated from murine or human lung tissue and co-cultured with murine CCL206 or human MRC5 fibroblasts, respectively, in Matrigel® (Corning Life Sciences B.V., Amsterdam, The Netherlands). EpCAM^+^ (CD31^-^/CD45^-^/CD326^+^) cells were isolated from mouse lung tissue (without the trachea) using the QuadroMACS™ Separator and antibody-bound magnetic beads (Miltenyi Biotec, Leiden, The Netherlands). EpCAM^+^ cells and fibroblasts were mixed 1:1 (20,000 cells each) and suspended in 100 μL of Matrigel prediluted 1:1 (v/v) with DMEM supplemented with 10% FBS. This mixture of cells was added to a 24-well Falcon® cell culture insert (Corning, USA) within a 24-well plate containing 400 μL of organoid media (DMEM/F-12 supplemented with 5% FBS, 1% penicillin/streptomycin, 1% glutamine, 1% amphotericin B, 0.025‰ EGF, 1% insulin-transferrin-selenium, and 1.75‰ bovine pituitary extract) underneath the insert in each well. Adult human donor tissue was isolated from histologically normal regions of lung tissue specimens obtained at University Medical Centre Groningen (Groningen, The Netherlands) from n = 7 patients (2 non-COPD and 5 COPD patients). Human lung tissues were incubated and homogenized overnight in an enzyme mixture at 4 °C; the EpCAM^+^ isolation process was similar to that described above for murine lung tissue. Organoids were cultured in a humidified atmosphere under 5% CO2/95% air at 37 °C and medium in the wells was refreshed every 2–3 days. To quantify the number of organoids, light microscopy at 20x magnification was used and organoids were counted manually. The diameter of the organoids (organoid size) was measured using NIS-Elements software with a light microscope.

For *in vitro* organoid experiments, organoids were continuously treated with control, 1.25% (1% for human organoids), 2.5% or 5% CSE and organoid culture medium was refreshed every other day. All information on the pharmacological compounds used in this study is provided in the Supplementary table 1.

### Immunofluorescence staining

The immunofluorescence staining assay for organoids was performed as described previously by our group with minor modifications^6,9,56^. Organoids were fixed in acetone diluted 1:1 (v/v) with methanol for 15 min at –20 °C. After fixation, one mL of PBS with 0.02 % sodium azide was added to the well underneath the insert. Organoids were kept at 4°C for one week after fixation. BSA media was added on top of the insert for blocking at room temperature (RT) for 2h. Afterwards, primary antibody incubation was performed in PBS buffer with 0.1% BSA and 0.1% Triton X-100 overnight at 4 °C. The next day, the organoids were washed with PBS for 30 min three times and secondary antibody incubation was performed for 2h at RT. After washing with PBS for 15 min, the organoids on the insert membrane were transferred to a glass slide with two drops of mounting medium containing DAPI (Abcam 104139, Cambridge, UK), and a coverslip was applied.

The slides were kept at 4 °C. Confocal images were acquired using a Leica SP8 microscope or a Leica DM4000B microscope. Images were obtained and analyzed with LASX (Leica) software (open resource, Leica Microsystems GmbH, Wetzlar, Germany).

### RNA extraction and RNA sequencing (RNA-seq) analysis

The EpCAM^+^ cells isolated from mice exposed to Air, CS, CS+misoprostol, or CS+iloprost were used to extract total RNA for RNA sequencing using NucleoSpin® RNA kit (Machery-Nagel, 740955, Germany) according to the manufacturer’s instructions. RNA concentrations and qualities were analyzed using Nanodrop spectrophotometer. An Illumina NovaSeq 6000 sequencer was used for RNA-seq data analysis by GenomeScan (https://www.genomescan.nl). The procedure included data quality control, adapter trimming, alignment of short reads and feature counting. Library preparation was checked by calculating ribosomal (and globin) content. Checks for possible sample and barcode contamination were performed and a set of standard quality metrics for the raw data set was determined using quality control tools (FstQC v0.34 and FastQA). Prior to alignment, the reads were trimmed for adapter sequences using Trimmomatic v0.30. To align the reads of each sample, the ensemble mouse reference GRCm38 (patch 6) was used. Data analyses following the RNA-seq studies were performed using the BioJupies platform (https://amp.pharm.mssm.edu/biojupies/)^57^. Gene expression in murine lung published data sets was obtained by downloading series matrix files from the NCBI GEO database GSE151674.

### Statistics analysis

All data are presented as mean ± SEM unless indicated otherwise. Unless stated otherwise, all data were assessed for statistical significance using two-tailed Student’s t-test or one-way ANOVA. The p-value indicating statistically significant differences between the mean/median values are defined as follows: *p < 0.05, **p < 0.01, ***p < 0.001, ****p < 0.0001. Statistical analyses were performed with GraphPad Prism 8 software.

## Acknowledgements

The authors would like to acknowledge the support from the Lung foundation Netherlands (Longfonds, grant 5.1.17.166) and China Scholarship Council (CSC201707720065).

## Author Contributions

Author X.W., R.G. conceptualized the project, analyzed and interpreted the data; R.G., L.E.M.K., and M.S. supervised the project; X.W., A.M., M.K., L.E.M.K., A.Ö.Y., and R.G. assisted designing the experiments; X.W., S.B., V.V., L.A.V., and A.D’M. performed the experiments; X.W., T.M.C., and M.A. prepared the figures; T.M.C., M.A., H.B.S., and A.Ö.Y. assisted in bioinformatics analysis; X.W., and R.G. wrote the manuscript; All authors reviewed and commented on the manuscript and agreed to the final version.

## Supplementary tables and figures

**Supplementary Table 1.**
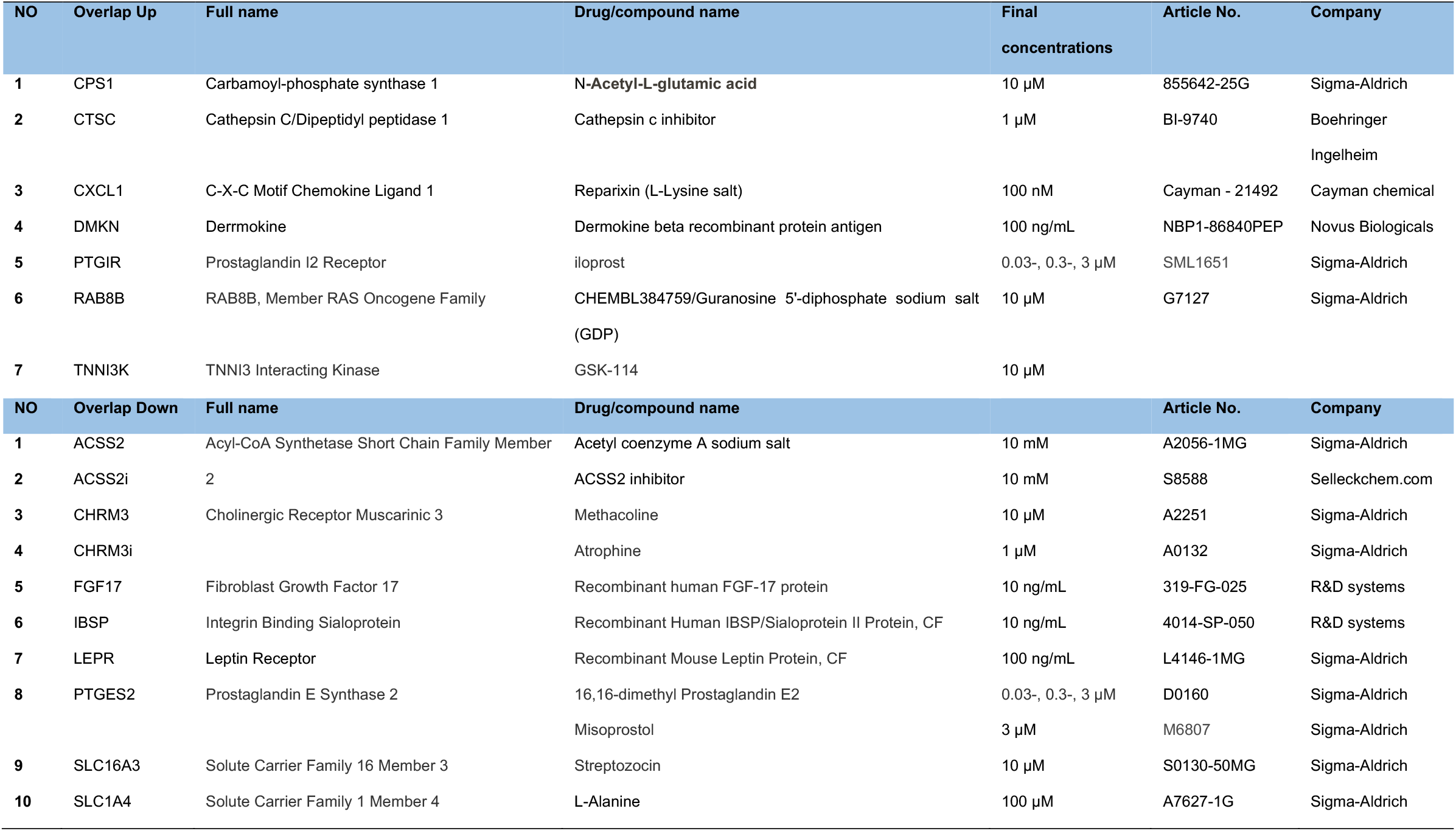
Information of final drug lists screened on organoid assay.

| NO | Overlap Up | Full name | Drug/compound name | Final concentrations | Article No. | Company |
| --- | --- | --- | --- | --- | --- | --- |
| 1 | CPS1 | Carbamoyl-phosphate synthase 1 | N-Acetyl-L-glutamic acid | 10 µM | 855642-25G | Sigma-Aldrich |
| 2 | CTSC | Cathepsin C/Dipeptidyl peptidase 1 | Cathepsin c inhibitor | 1 µM | BI-9740 | Boehringer<br>Ingelheim |
| 3 | CXCL1 | C-X-C Motif Chemokine Ligand 1 | Reparixin (L-Lysine salt) | 100 nM | Cayman - 21492 | Cayman chemical |
| 4 | DMKN | Dermokine | Dermokine beta recombinant protein antigen | 100 ng/mL | NBP1-86840PEP | Novus Biologicals |
| 5 | PTGIR | Prostaglandin I2 Receptor | iloprost | 0.03-, 0.3-, 3 µM | SML1651 | Sigma-Aldrich |
| 6 | RAB8B | RAB8B, Member RAS Oncogene Family | CHEMBL384759/Guanosine 5'-diphosphate sodium salt (GDP) | 10 µM | G7127 | Sigma-Aldrich |
| 7 | TNNI3K | TNNI3 Interacting Kinase | GSK-114 | 10 µM |  |  |
| NO | Overlap Down | Full name | Drug/compound name |  | Article No. | Company |
| 1 | ACSS2 | Acyl-CoA Synthetase Short Chain Family Member | Acetyl coenzyme A sodium salt | 10 mM | A2056-1MG | Sigma-Aldrich |
| 2 | ACSS2i | 2 | ACSS2 inhibitor | 10 mM | S8588 | Selleckchem.com |
| 3 | CHRM3 | Cholinergic Receptor Muscarinic 3 | Methacoline | 10 µM | A2251 | Sigma-Aldrich |
| 4 | CHRM3i |  | Atrophine | 1 µM | A0132 | Sigma-Aldrich |
| 5 | FGF17 | Fibroblast Growth Factor 17 | Recombinant human FGF-17 protein | 10 ng/mL | 319-FG-025 | R&D systems |
| 6 | IBSP | Integrin Binding Sialoprotein | Recombinant Human IBSP/Sialoprotein II Protein, CF | 10 ng/mL | 4014-SP-050 | R&D systems |
| 7 | LEPR | Leptin Receptor | Recombinant Mouse Leptin Protein, CF | 100 ng/mL | L4146-1MG | Sigma-Aldrich |
| 8 | PTGES2 | Prostaglandin E Synthase 2 | 16,16-dimethyl Prostaglandin E2 | 0.03-, 0.3-, 3 µM | D0160 | Sigma-Aldrich |
|  |  |  | Misoprostol | 3 µM | M6807 | Sigma-Aldrich |
| 9 | SLC16A3 | Solute Carrier Family 16 Member 3 | Streptozocin | 10 µM | S0130-50MG | Sigma-Aldrich |
| 10 | SLC1A4 | Solute Carrier Family 1 Member 4 | L-Alanine | 100 µM | A7627-1G | Sigma-Aldrich |

**Supplementary Table 2.** Information of compounds used in organoid assay.

| Name | Final concentrations | Article No. | Company |
| --- | --- | --- | --- |
| Rolipram | 1-, 10 µM | R6520 | Sigma-Aldrich |
| Budesonide | 1-, 10-, 100 nM | B7777 | Sigma-Aldrich |
| Cholera toxin | 0.1 mg/mL | C8052 | Sigma-Aldrich |
| (R) -Butaprost (EP2 analogue) | 30 µM | B6309 | Sigma-Aldrich |
| EP4 analogue* | 30 µM | CAY10598 | Cayman chemical |
| Forskolin | 10 µM | F6886 | Sigma-Aldrich |
| IBMX (3-Isobutyl-1-methylxanthine) | 100 µM | Obtained from AppliChem (Germany) |  |
| db-cAMP (Bucladesine) | 100 µM | D0627 | Sigma-Aldrich |
| Olodaterol | 100 nM | Obtained from AppliChem (Germany) |  |

\*EP4 analouge: 5-[(3S)-3-hydroxy-4-phenyl-1-buten-1-yl]1-[6-(2H-tetrazol-5R-yl)hexyl]-2-pyrrolidinone

**Supplementary figure 1.**
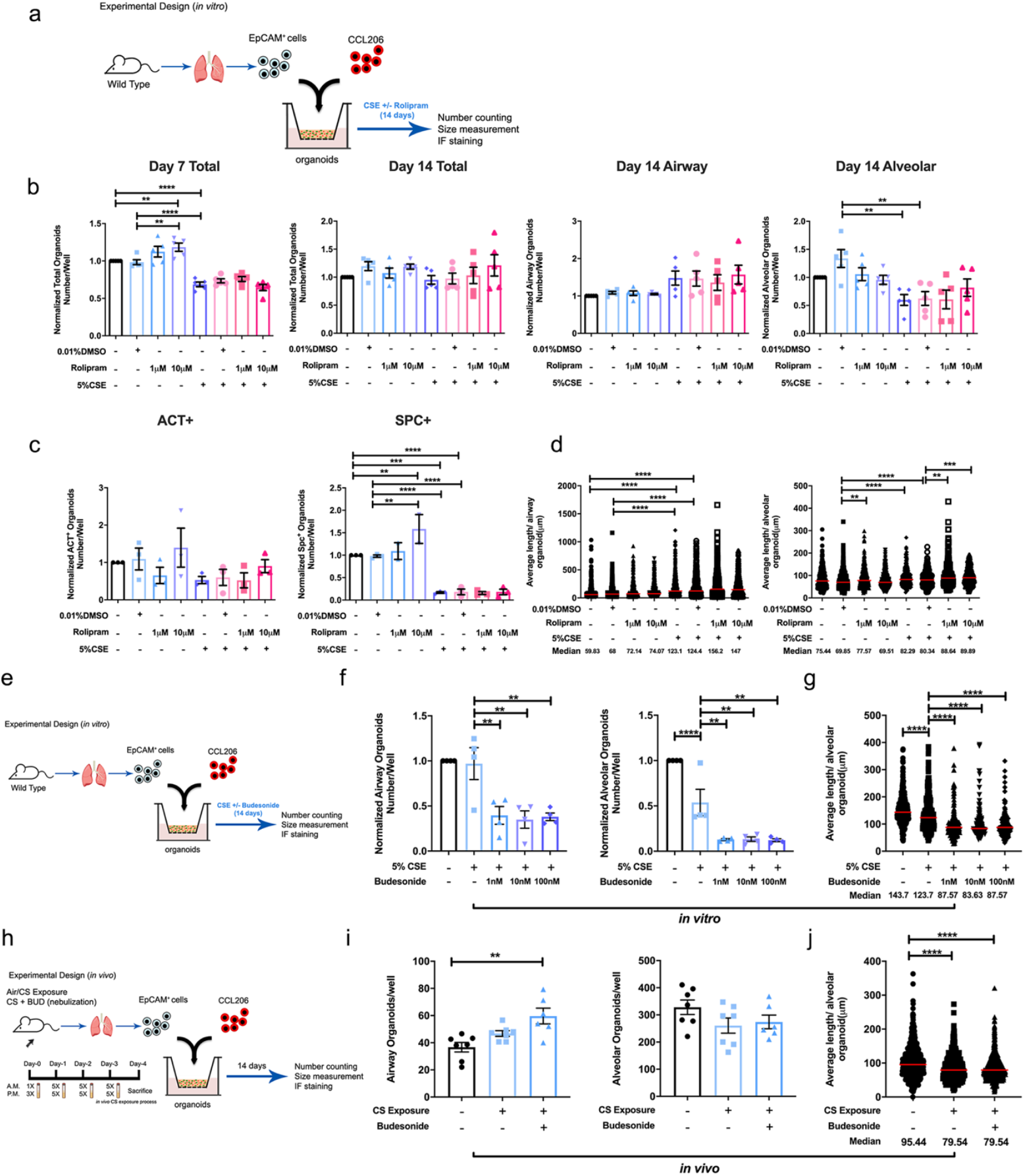
Effect of Rolipram and Budesonide on lung organoid formation. **a** Schematic of *in vitro* experimental design. **b** Quantification of normalized number of total organoids (day 7), total organoids (day 14), airway type organoids (day 14), alveolar type organoids (day 14) treated with 5% CSE ± rolipram (0-, 1-, 10μM). **c** Quantification of normalized ACT^+^ and SPC^+^ organoid numbers treated with 5% CSE ± rolipram (0-, 1-, 10μM) at Day 14. **d** Quantification of average length (diameter) of airway and alveolar type organoids treated with 5% CSE ± rolipram (0-, 1-, 10μM) measured on day 14. N = 5 experiments, n > 503 organoids/group. Data are presented as scatter plots with medians. **e** Schematic of *in vitro* experimental design. **f** Quantification of normalized number of airway and alveolar type organoids treated with 5% CSE ± Budesonide (0-, 1-, 10-, 100 nM) measured on day 14. **g** Quantification of average length (diameter) of alveolar type organoids (median value) treated with 5% CSE ± Budesonide (0-, 1-, 10-, 100 nM) measured on day 14. N = 4 experiments, n > 165 organoids/group. Data are presented as scatter plots with medians. **h** Schematic of *in vivo* experimental design. **i** Number of airway and alveolar type organoids from co-culture of CCL-206 fibroblasts and EpCAM^+^ cells (isolated from air-exposed, CS-exposed, and CS-exposed + Budesonide nebulized mice) quantified on day 14. **j** Quantification of average length (diameter) of alveolar organoids (median value) from co-culture of CCL-206 fibroblasts and EpCAM^+^ cells (isolated from air-exposed, CS-exposed, and CS-exposed + Budesonide nebulized mice) on day 14. N = 6 - 7 experiments, n > 671 organoids/group. Data are presented as scatter plots with medians. **p < 0.05, **p < 0.01, ***p < 0.001, ****p < 0.0001

**Supplementary figure 2.**
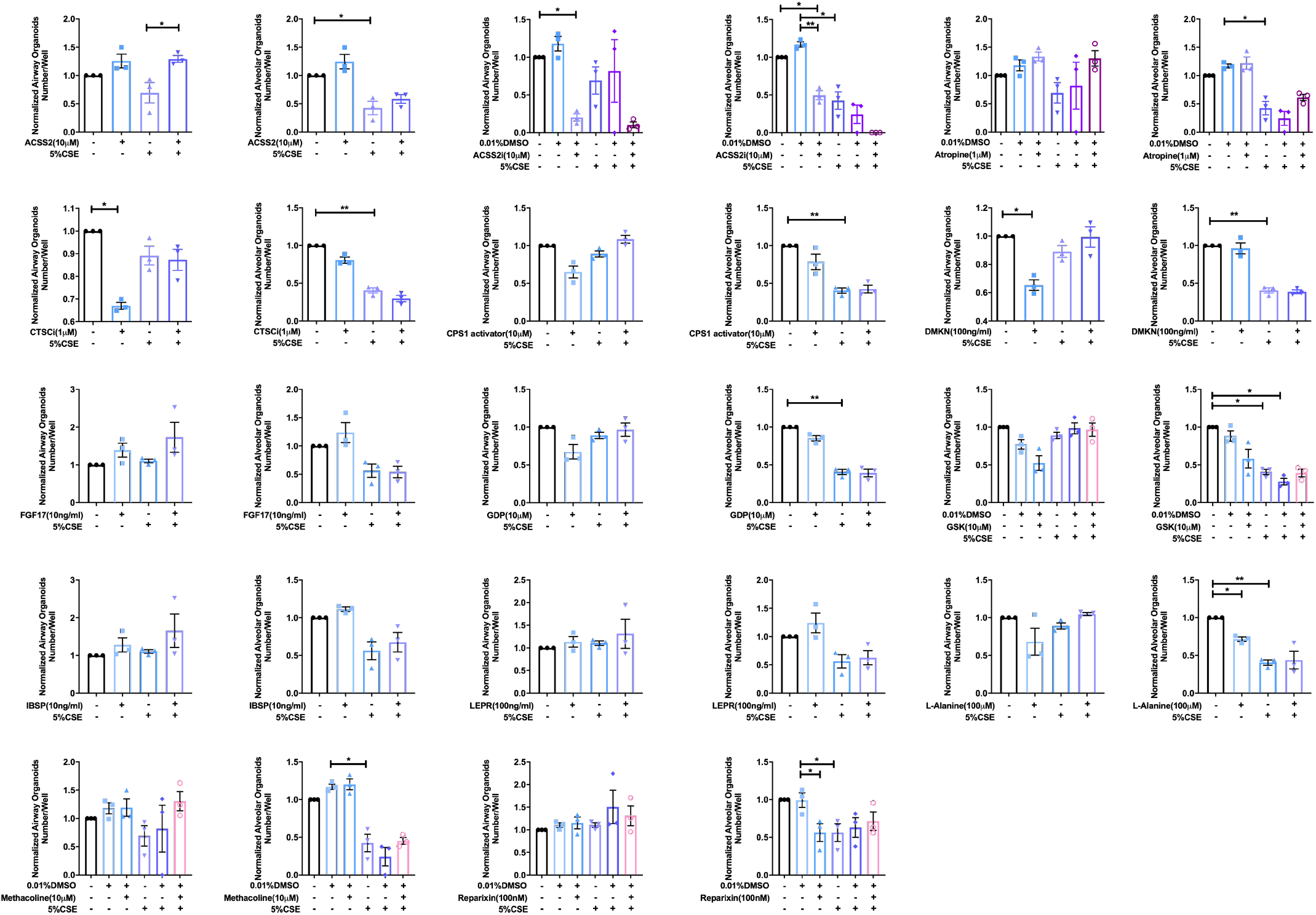
Effects of 15 drugs candidates of interest (including an additional agonist and antagonist for one gene target) on the normalized number of airway and alveolar type lung organoids in the presence and absence of 5% CSE. Data are presented as median ± SEM. **p < 0.05, **p < 0.01, ***p < 0.001, ****p < 0.0001.

**Supplementary figure 3.**
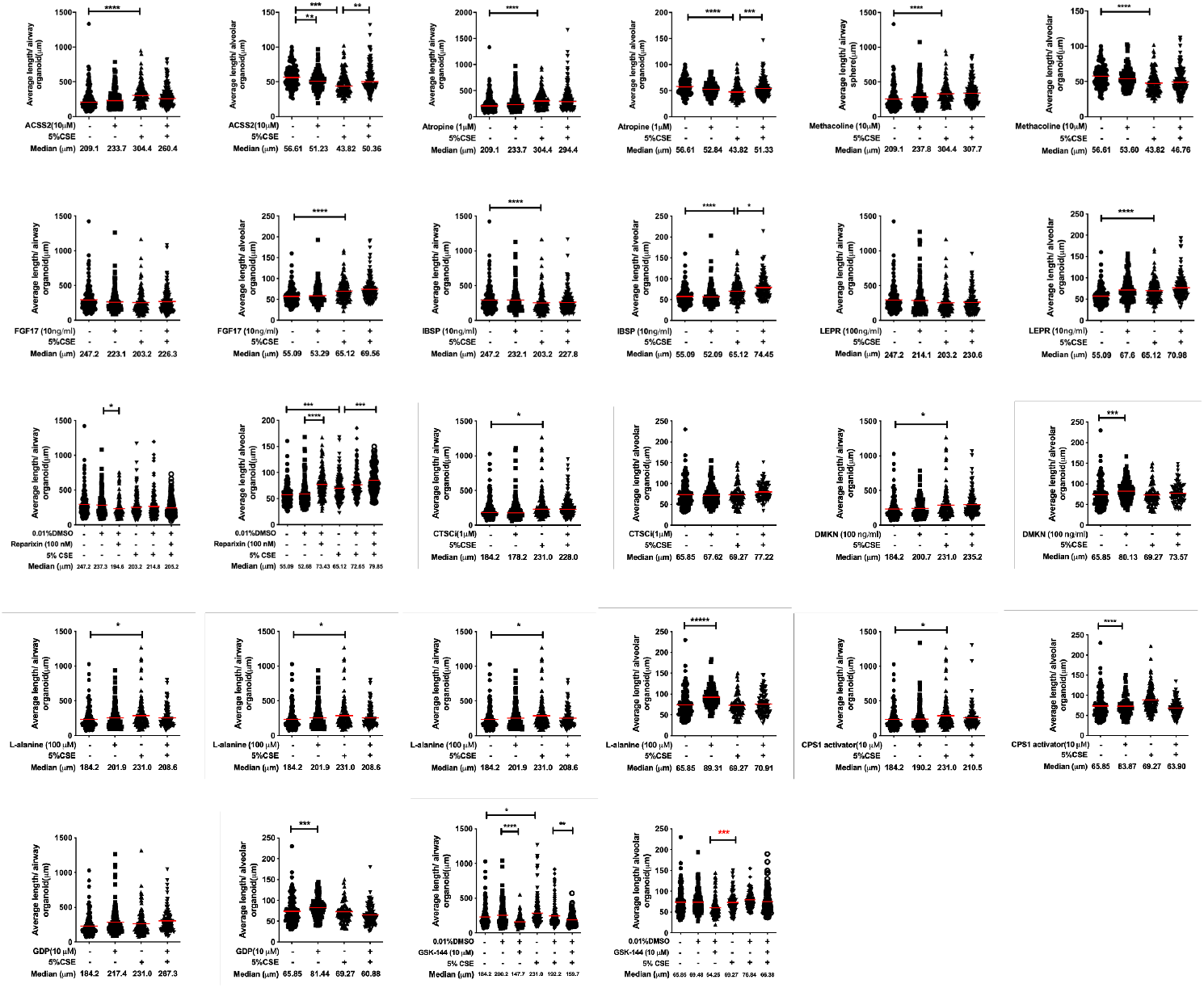
Effects of 15 drugs candidates of interest (including an additional agonist and antagonist for one gene target) on the size of airway and alveolar type lung organoids in the presence and absence of 5% CSE. Data are presented as median ± SEM. **p < 0.05, **p < 0.01, ***p < 0.001, ****p < 0.0001.

**Supplementary figure 4.**
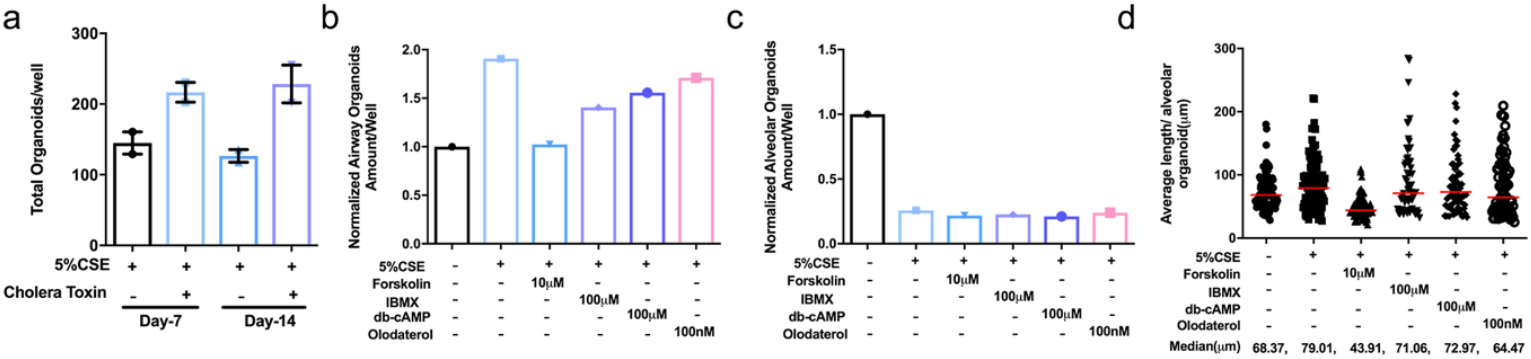
Compounds related to cAMP signaling pathways tested on lung organoid assay. **a** Quantification of total organoids treated with 5% CSE + cholera toxin at different time points. **b** Quantification of normalized airway type organoids treated with 5% CSE ± Forskolin (10 µM), IBMX (100 µM), db-cAMP (100 µM), and olodaterol (100 nM). **c** Quantification of normalized alveolar type organoids treated with 5% CSE ± Forskolin (10 µM), IBMX (100 µM), db-cAMP (100 µM), and olodaterol (100 nM). **d** Quantification of average length (diameter) of alveolar type organoids (median value) treated with 5% CSE ± Forskolin (10 µM), IBMX (100 µM), db-cAMP (100 µM), and olodaterol (100 nM). Data are presented as median ± SEM. **p < 0.05, **p < 0.01, ***p < 0.001, ****p< 0.0001

**Supplementary figure 5.**
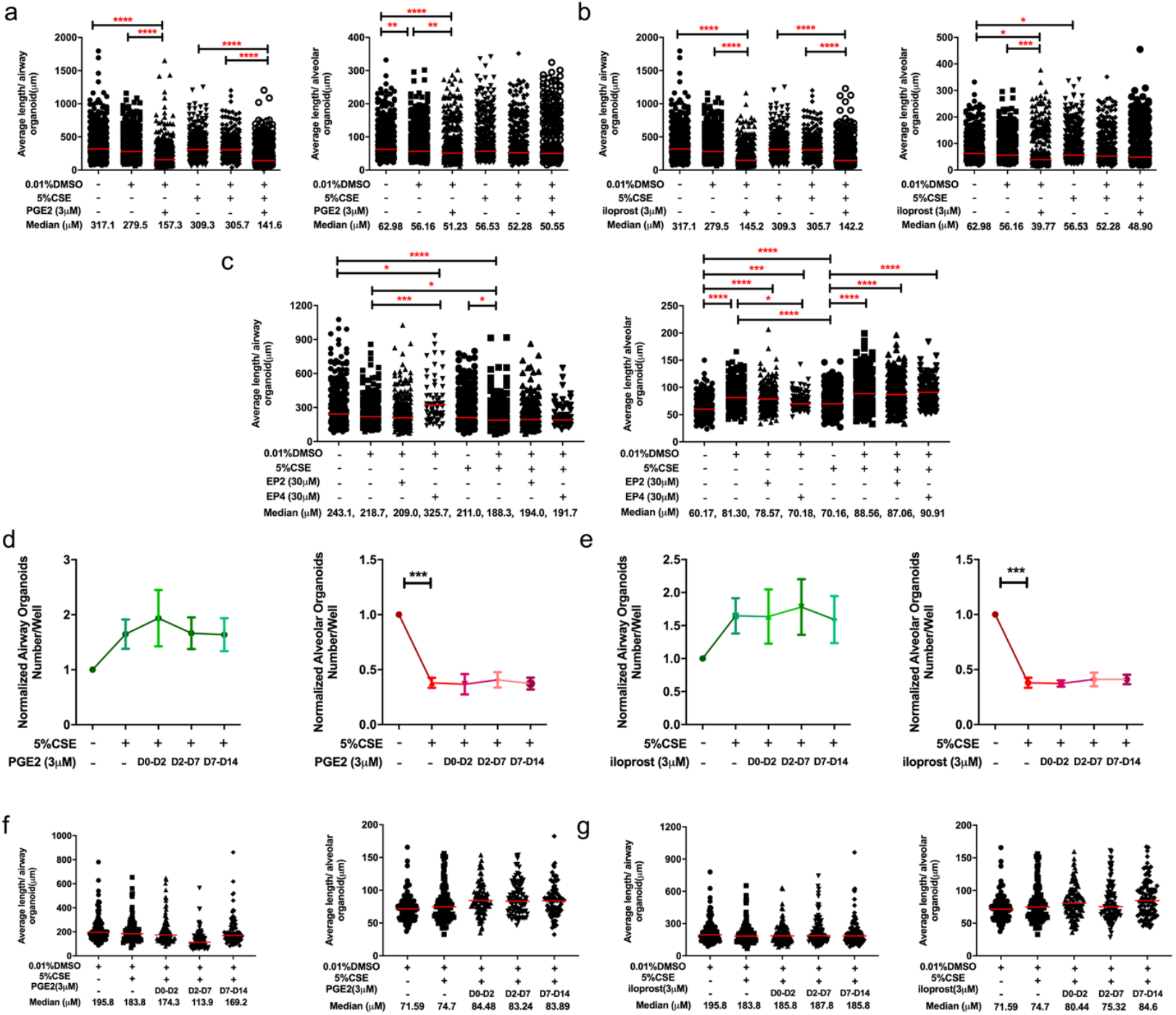
Quantification of lung organoid numbers and sizes in the study of PGE2/PGI2/EP2/EP4. **a-b** Quantification of average length (diameter) of organoids (median value) of airway and alveolar type organoids treated with 5% CSE ± PGE2 agonist (a)/iloprost (b) measured on day 14. N = 5 experiments, n > 334 organoids/group. Data are presented as median ± SEM. **c** Quantification of average length (diameter) of airway and alveolar type organoids (median value) treated with 5% CSE ± EP2/EP4 agonists measured on day 14. N = 5 experiments, n > 264 organoids/group. Data are presented as median ± SEM. **d-e** Quantification of normalized number of airway and alveolar type organoids treated with vehicle control, 5% CSE ± PGE2 agonist (d)/ iloprost (e) from day 0-2, day 2-7, or day 7-14. **f-g** Quantification of average length (diameter) of airway and alveolar type organoids (median value) treated with 5% CSE ± PGE2 agonist (f)/ iloprost (g) from day 0-2, day 2-7, and day 7-14. N = 2 experiments, n > 85 organoids/group. Data are presented as median ± SEM.

**Supplementary figure 6.**
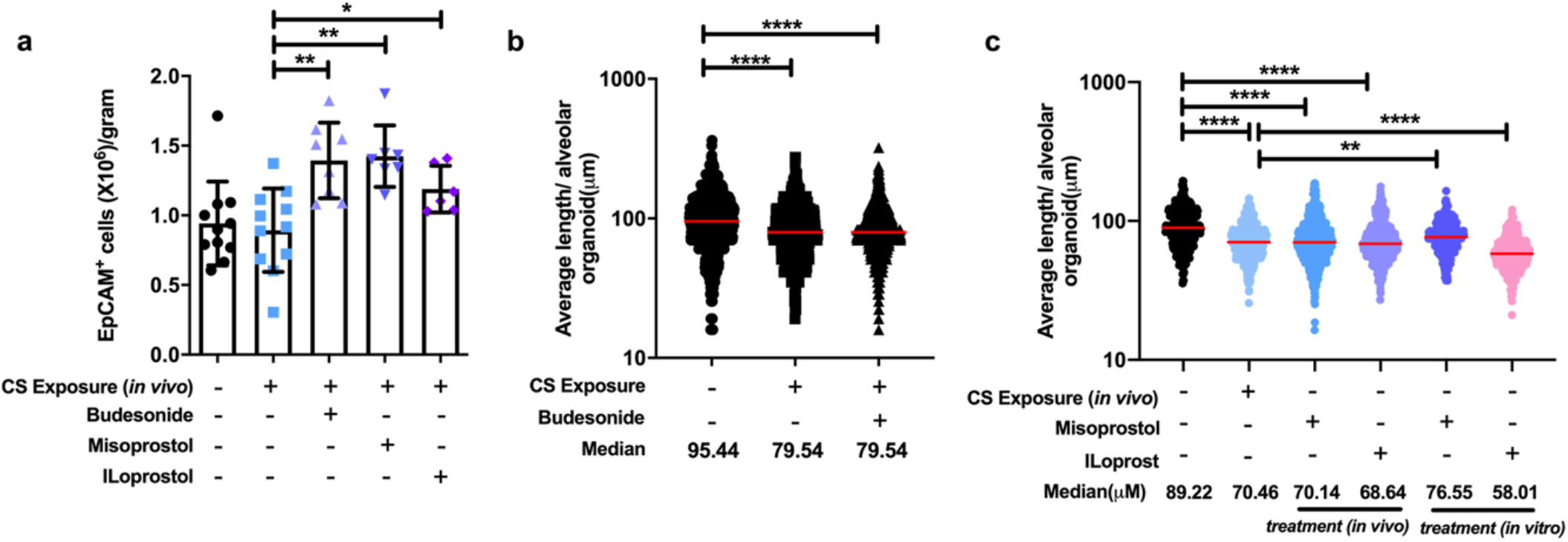
Quantification analysis of lung organoid assay in the *in vivo* study of PGE2/PGI2/Budesonide. **a** Yield efficiency of EpCAM^+^ cells from mice with different treatments. **b** Quantification of average length of alveolar type organoids co-cultured from CCL-206 and EpCAM^+^ cells isolated from air- (control) or CS-exposed mice with or without *in vivo* treatment with misoprostol or iloprost (i.p injection). N = 4-8 experiments, n = 400 organoids/group. **c** Quantification of average length of alveolar type organoids co-cultured from CCL-206 and EpCAM^+^ cells isolated from air- or CS-exposed mice. Organoids were treated *in vitro* with misoprostol or iloprost for 14 days. N = 4-8 experiments, n = 400 organoids/group. Data are presented as median ± SEM. **p < 0.05, **p < 0.01, ***p < 0.001, ****p < 0.0001.

